# HLA-DQA1*05 is associated with the development of antibodies to anti-TNF therapy

**DOI:** 10.1101/410035

**Authors:** Aleksejs Sazonovs, Nick Kennedy, Loukas Moutsianas, Graham A. Heap, Daniel L. Rice, Mark Reppell, Claire Bewshea, Gareth Walker, Mandy H. Perry, Timothy J. McDonald, Charlie Lees, Fraser Cummings, Miles Parkes, John Mansfield, Jeffrey C. Barrett, Dermot McGovern, James Goodhand, Carl A. Anderson, Tariq Ahmad, PANTS consortium

## Abstract

**Background:** Anti-tumour necrosis factor (anti-TNF) therapies are the most widely used biologic therapies for treating immune-mediated diseases. Their efficacy is significantly reduced by the development of anti-drug antibodies which can lead to treatment failure and adverse reactions. The biological mechanisms underlying antibody development are unknown but the ability to identify subjects at higher risk would have significant clinical benefits.

**Methods:** The PANTS cohort consists of Crohn’s disease patients recruited prior to first administration of anti-TNF, with serial measurements of anti-drug antibody titres. We performed a genome-wide association study across 1240 individuals from this cohort to identify genetic variants associated with anti-drug antibody development.

**Findings:** The Human Leukocyte Antigen allele, HLA-DQA1*05, carried by approximately 40% of Europeans, significantly increased the rate of anti-drug antibody development (hazard ratio [HR], 1.90; 95% confidence interval [CI], 1.60 to 2.25; P=5.88×10^-13^). This association was consistent for patients treated with adalimumab (HR, 1.89; 95% CI, 1.32 to 2.70) and infliximab (HR, 1.92; 95% CI, 1.57 to 2.33), and for patients treated with mono-(HR, 1.75; 95% CI, 1.37 to 2.22) or combination therapy with immunomodulators (HR, 2.0; 95% CI, 1.57 to 2.58).

**Interpretation:** HLA-DQA1*05 is significantly associated with an increased rate of anti-drug antibody formation in patients with Crohn’s disease treated with infliximab and adalimumab. Pre-treatment HLA-DQA1*05 genetic testing may help personalise the choice of anti-TNF therapy and allow the targeted use of immunomodulator therapy to minimise risk and maximise response.

## Introduction

Biological therapies, commonly known as biologics, are typically large and complex proteins manufactured in, or derived from, living sources. Biologics have transformed the management of immune-mediated diseases and in 2017 accounted for a global expenditure in excess of $100 billion.^1^ However, repeated exposure stimulates the formation of anti-drug antibodies – the most frequently cited cause of treatment failure, hypersensitivity reactions, and treatment discontinuation.^2^

The anti-tumour necrosis factor antibodies (anti-TNFs), infliximab and adalimumab, are the most commonly prescribed biologics for treating inflammatory bowel disease (IBD)^3^. The risk of development of anti-drug antibodies, referred to as immunogenicity, is higher for patients treated with infliximab (a murine-human chimeric monoclonal antibody) than adalimumab (a fully humanised monoclonal antibody).^2^ Combination immunomodulator therapy reduces immunogenicity to both adalimumab and infliximab and for infliximab improves treatment outcomes.^4,5^ Despite these benefits, many patients are still treated with anti-TNF monotherapy because of concerns about the increased risk of adverse drug reactions, opportunistic infections, and malignancies associated with combination therapy.^6–10^

Knowledge of the cellular and molecular mechanisms underpinning failure of immune tolerance to biologics is limited. Retrospective candidate gene studies have suggested variants in *FCGR3A*^*11*^ and *HLA-DRB1*^*12,13*^ increase susceptibility to immunogenicity to anti-TNF therapy. However, these associations did not achieve genome-wide significance and are yet to be independently replicated.

Here we report the first genetic locus robustly associated with immunogenicity to anti-TNF therapies using data from the Personalising Anti-TNF Therapy in Crohn’s disease (PANTS) study.

## Methods

120 hospitals from across the United Kingdom recruited 1610 patients to the PANTS study (NHS REC 12/SW/0323; IRAS 115956; ClinicalTrials.gov identifier: NCT03088449) at the time of first anti-TNF exposure. To be included in the study, patients had to be aged 6 years or over with active luminal Crohn’s disease involving the colon and/or small intestine where the primary indication for anti-TNF treatment was not fistulising disease. Choice of anti-TNF drug and use of concomitant immunomodulator therapy was at the discretion of the treating physician as part of usual care. Patients were initially studied for 12 months, or until drug withdrawal by the treating physician. In the first year, study visits were scheduled at first dose, post-induction (weeks 12-14), weeks 30, 54 and at treatment failure. For infliximab-treated patients, additional visits occurred at each infusion. After 12 months patients were invited to continue follow-up for a further two years. Patients who declined the two-year extension were censored at the time of study exit.

At each visit, serum infliximab or adalimumab drug and anti-drug antibody levels were analysed on the Dynex (Chantilly Virginia, USA) DS2 automated enzyme linked immunosorbent assay (ELISA) platform, using the Immundiagnostik (Immundiagnostik AG, Bensheim, Germany) IDKmonitor^®^ drug (K9655 infliximab drug level and K9657 adalimumab drug level) and total antibody ELISA assays (K9654 infliximab total anti-drug antibody and K9651 adalimumab total anti-drug antibody). Based on our own independent assay experiments,^14^ we defined immunogenicity as an anti-drug antibody concentration ≥10 AU/mL, irrespective of drug concentration.

### Genotyping, quality control and genotype imputation

Genotyping of DNA from 1524 patients was undertaken using the Illumina CoreExome microarray, with genotype calls made using optiCall.^15^ Pre-imputation quality control (QC) of samples and single nucleotide polymorphisms (SNPs) was performed as previously described.^16^ We excluded individuals of non-European ancestry, one individual from related pairs (defined as a pi-hat>0.1875), and individuals with an outlying number of missing or heterozygous genotypes. Variants with a genotype call rate of <95% were also excluded.

SNP genotypes were imputed via the Sanger Imputation Service using the Haplotype Reference Consortium (HRC) panel.^17^ Following imputation we ran additional variant and sample QC procedures (appendix methods).

HLA imputation was carried out using the HIBAG package^18^ in R, using pre-fit classifiers trained specifically for the CoreExome genotyping microarray on individuals of European ancestry. HLA types were imputed at 2-, and 4-digit resolution for the following loci: *HLA-A, HLA-C, HLA-B, HLA-DRB1, HLA-DQA1, HLA-DQB1*, and *HLA-DPB1*. 4-digit resolution separates alleles which differ in nucleotide positions that determine amino acid sequences, whereas the lower, 2-digit resolution separates allele families. In addition, we obtained amino acid sequences for all the imputed HLA alleles, using the IPD-IMGT/HLA database.^19^ Following the recommended best practices for HIBAG,^18^ HLA-allele and amino acid calls with posterior probability <0.5 were set to missing for the given individual. HIBAG’s imputation accuracy is very high in Europeans, above 94% across all HLA loci and above 99% for the *HLA-DQA1* locus specifically.^20^ Furthermore, we validated the imputation accuracy for HLA-DQA1*05 at 2D resolution using typed HLA data (appendix table S1).

### Genome-wide association analyses

Cox proportional hazards regression was used to test genetic variants and for association with time to anti-drug antibody development, assuming an additive genetic effect. Patients who did not develop immunogenicity during the study were censored at the point of last observation. The association analyses were performed using a likelihood ratio test with sex, drug type (infliximab or adalimumab), immunomodulator use, and the first principal component as covariates (appendix table S2), with the SurvivalGWAS_SV software.^21^ Kaplan–Meier estimators were produced using the Lifelines package^22^ for Python.

The Akaike information criterion (AIC), calculated using the ‘survival’ R package,^23^ was used for model comparison across non-nested models; this estimates the goodness of fit while penalising for the number of parameters to prevent overfitting. The fixed effects Q statistic, calculated using the ‘metafor’ R package,^24^ was used to perform tests of heterogeneity of effect; this test is an extension of Cochran’s Q-test and examines whether the observed effect size variability is larger than expected by chance.

### Replication cohort

We assembled an independent cohort to replicate significant findings from the discovery cohort. This comprised 107 Crohn’s disease, 64 ulcerative colitis, and 7 IBD type-unclassified patients, all with cross-sectional drug and antibody levels measured as part of routine clinical practice at the Exeter laboratory. The samples were genotyped using either the Illumina CoreExome array (N=164)^16^ or the Affymetrix 500k array (N=14).^25^ We repeated the same quality control procedures and imputation (genotype and HLA) undertaken on the PANTS discovery cohort. We applied the same statistical model used for the discovery analysis, with IBD disease type as an additional covariate.

## Results

Of the 1610 patients recruited to the PANTS study, 1524 were genotyped, and 1323 of these passed genetic QC, ancestry, and relatedness analyses. Drug and antibody level data were available for 1240 of these patients (appendix figure S2, appendix table S3), of which 44% developed anti-drug antibodies within the first 12 months (95% CI, 0.41 to 0.48), 62% within 36 months (95% CI, 0.57 to 0.67).

After correcting for immunomodulator use, the rate of immunogenicity was greater in patients treated with infliximab than adalimumab (HR, 3.21; 95% CI, 2.61 to 3.95; P<1.18×10^-28^). In a model including drug-type as a covariate, rate of immunogenicity was greater in patients treated with anti-TNF monotherapy (N=544) compared to combination therapy with immunomodulators (N=696), (HR, 2.30; 95% CI, 1.94 to 2.75; P<6.10×10^-21^).

### Genetic variants in the HLA region are associated with time to immunogenicity

We identified a genome-wide significant association with time to development of immunogenicity on chromosome 6 (figure 1, appendix figures S3 and S4), with the most associated SNP, rs2097432 (b38_pos: 6:32622994; HR, 1.70; 95% CI, 1.48 to 1.94; P=4.24×10^-13^), falling within the major histocompatibility complex (MHC) region (appendix figures S3 and S4). We replicated this association in an independent cohort (HR, 1.69; 95% CI, 1.26 to 2.28; P=8.80×10^-4^). A variant on chromosome 11, rs12721026 (b38_pos: 11:116835452; HR, 0.46; 95% CI, 0.33 to 0.63; P=4.76×10^-8^), also reached genome-wide significance in our discovery analysis, though the association was not replicated in our independent cohort (HR, 0.85; 95% CI, 0.49 to 1.44; P=0.51).

**Figure 1:**
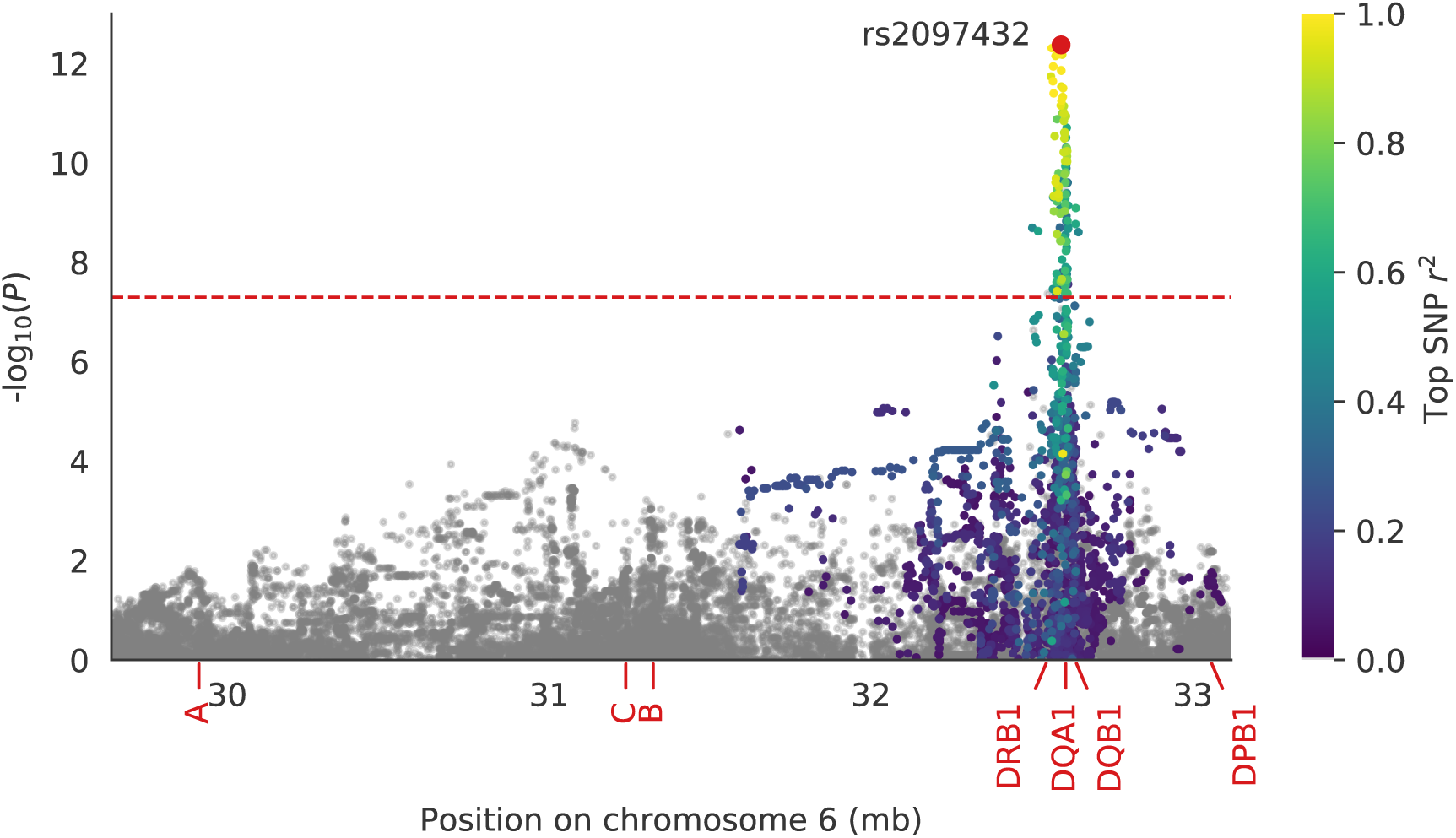
Regional plot of the association results with the MHC region on chromosome 6. Midpoint positions of the HLA alleles across the MHC region are shown in red on the x-axis. SNPs that passed the genome-wide significance threshold (P=5×10^-8^) are shown above the red horizontal dashed line with the most significant SNP in red. SNPs correlated with the lead SNP (r^2^>0.05) are color-coded from purple to yellow. Pairwise genotype correlation (r^2^) between SNPs was calculated using genotype data from the non-Finnish European population of 1KGP.^26^

### Fine-mapping of the signal in the HLA region

At the 2-digit level (HLA allele group), HLA-DQA1*05 was the only allele to reach genome-wide significance (HR, 1.58; 95% CI, 1.39 to 1.80; P=1.94×10^-11^). At the 4-digit level, no single allele reached genome-wide significance. The two most common HLA-DQA1*05 subtype alleles, HLA-DQA1*05:01 (HR, 1.57; 95% CI, 1.33 to 1.85; P=4.24×10^-7^) and HLA-DQA1*05:05 (HR, 1.48; 95% CI, 1.24 to 1.78; P=5.54×10^-5^), had similar effects on time to immunogenicity. However, a model using the HLA-DQA1*05 allele group was a better fit than using our two most common subtype alleles (AIC_05_=6659.07 versus AIC_05:01&05:05_=6659.50).

Carriage of two HLA-DQA1*05 alleles does not appear to substantially shorten the time to immunogenicity compared to carriage of one risk allele (figure 2). To formally assess this, we compared the fit of both the additive and dominant models for HLA-DQA1*05 and found that a dominant model led to a better fit (AIC_DOM_=6652.12 vs AIC_ADD_=6659.07), and gave the strongest association signal for HLA-DQA1*05 (HR, 1.90; 95% CI, 1.60 to 2.25; P=5.88×10^-13^). We also looked for non-additive effects across all other HLA alleles, but the model assuming a dominant effect for HLA-DQA1*05 remained the best fit to the data (appendix table S4).

**Figure 2:**
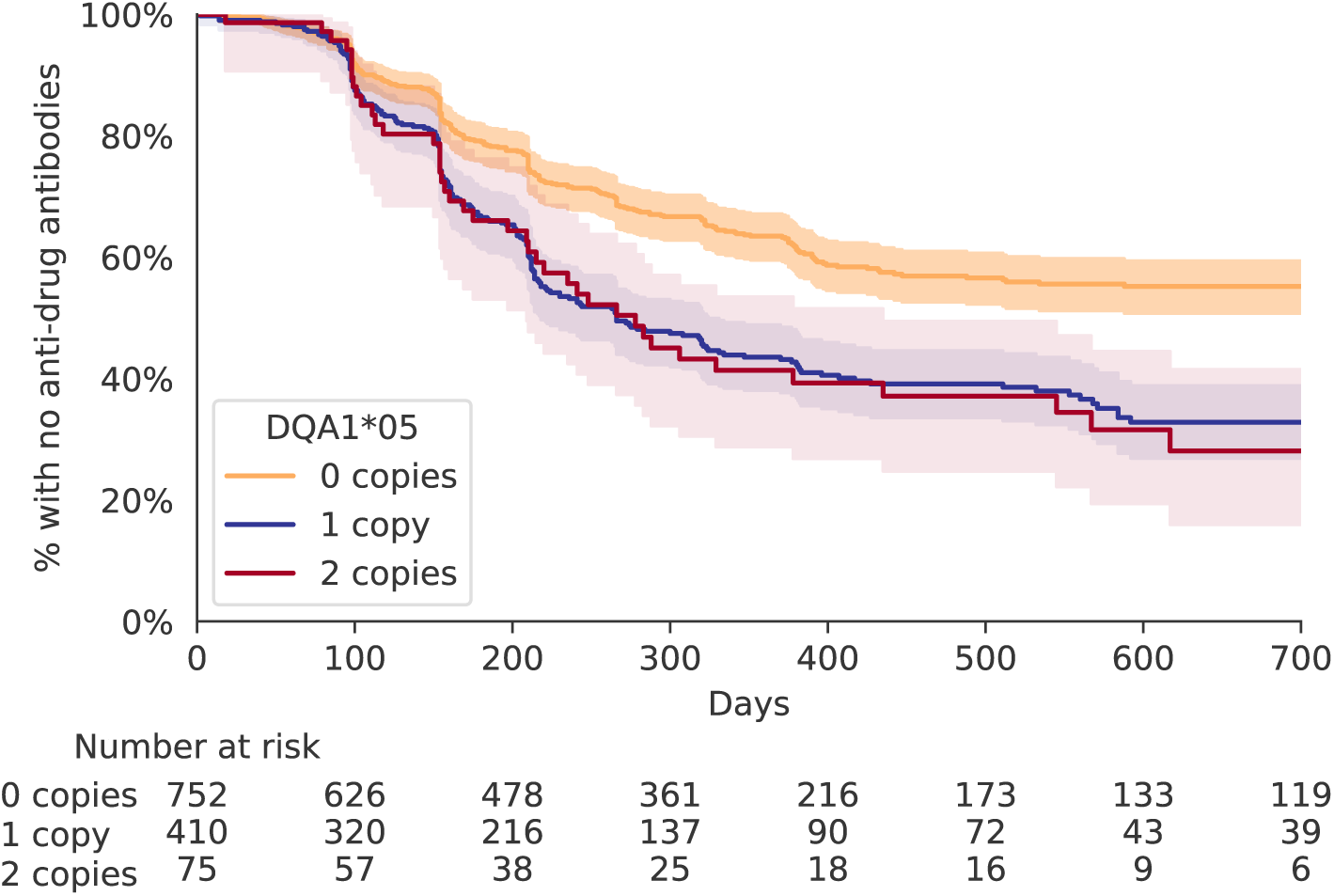
Kaplan–Meier estimator showing the rate of anti-drug antibody development, stratified by the number of HLA-DQA1*05 alleles carried. Orange, blue and red indicate 0, 1 and 2 copies of DQA1*05 allele, respectively. Carriers of one or two copies of the allele have a similar rate of immunogenicity development, and a dominant model is a better fit for the data than an additive model (AIC_DOM_=6652.12 vs AIC_ADD_=6659.07). X-axis truncated at 700 days, due to the low number of observations for longer time periods.

After conditioning on HLA-DQA1*05 we did not identify any secondary signals of association with time to immunogenicity within the MHC region (appendix figure S6). The HLA-DQA1*05 association was confirmed in our replication cohort (HR, 2.00; 95% CI, 1.35 to 2.98; P=6.60×10^-4^), again with a better fit for the dominant model (AIC_DOM_=942.51 vs AIC_ADD_=944.81). The association was robust to the antibody titre used to define immunogenicity (appendix figure S5).

We next looked for associations between specific amino acid residues within the classical HLA loci and time to immunogenicity.^27^ Isoleucine at position 107 (AA-107-ile) and lysine at position 175 (AA-175-lys) of the mature protein sequence are perfectly tagging the HLA-DQA1*05 alleles (HLA-DQA1*05:01, HLA-DQA1*05:03, HLA-DQA1*05:05, HLA-DQA1*05:09), so their effect is virtually indistinguishable from that of HLA-DQA1*05 (figure 3). Any differences between the models are due to minor inconsistencies in the imputation results for HLA-DQA1*05 alleles at different resolutions. No other amino acid residues showed comparable association signals.

**Figure 3:**
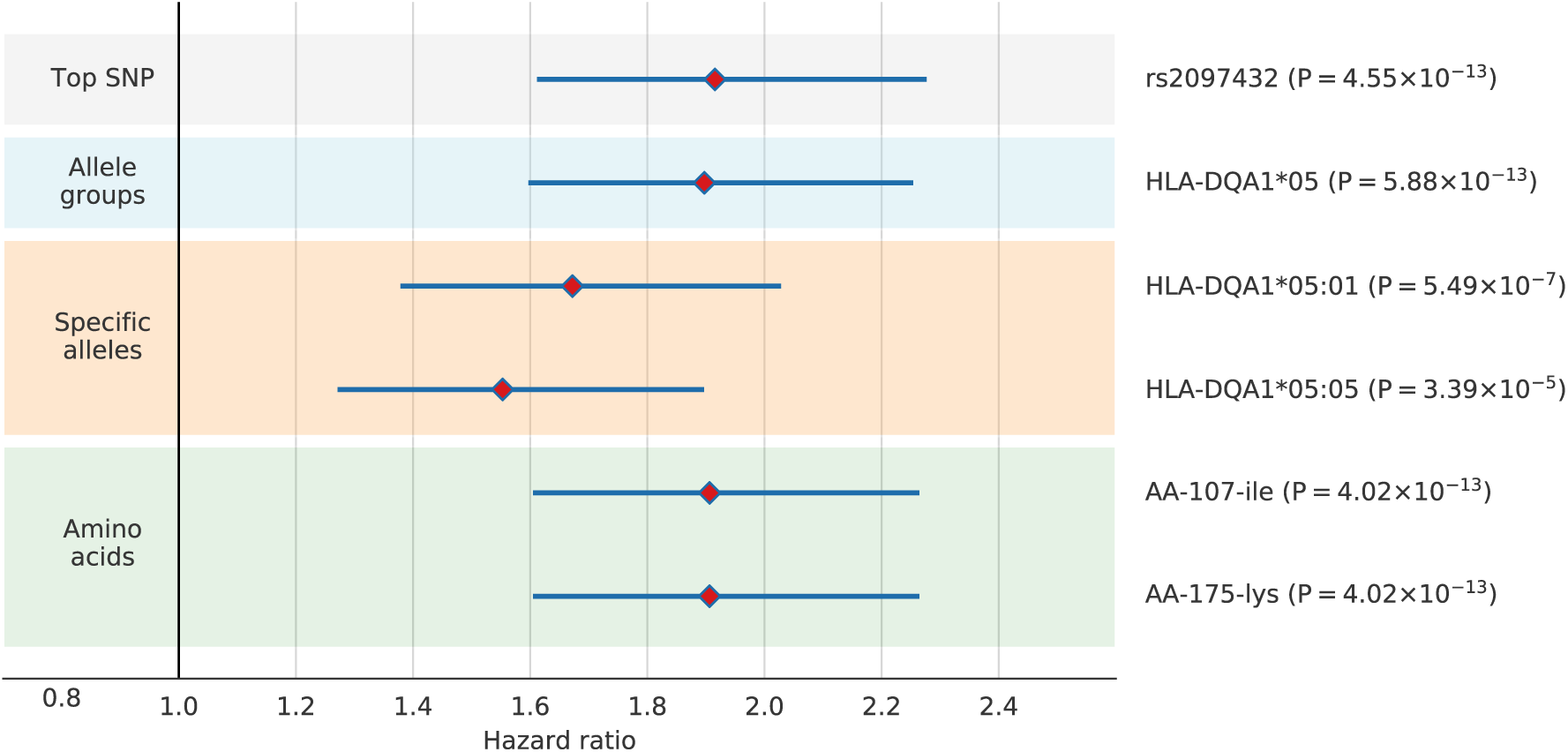
Effect sizes of the most strongly associated SNP, HLA alleles and amino acids on time to immunogenicity. Blue lines represent 95% CIs. Association test P-values are shown in parentheses.

### The effect of HLA-DQA1*05 across drug and treatment regimes

While immunogenicity rates were lower with adalimumab versus infliximab treated patients, the effect of HLA-DQA1*05 carriage was the same for the two therapies (P_het_=0.91) (figure 4). We also found no evidence for heterogeneity of the HLA-DQA1*05 effect on immunogenicity (P_het_=0.23) between patients treated with the infliximab originator, Remicade (Merck Sharp & Dohme Corp), and its biosimilar CT-P13 (Celltrion, South Korea; branded as Remsima, Napp UK, and Inflectra, Pfizer, USA) (appendix figure S7). These findings support the previously reported similar antigenic profile for infliximab originator and CT-P13.^28^ Likewise, the effect of HLA-DQA1*05 carriage on immunogenicity was consistent for individuals on mono-versus combination therapy with immunomodulators (P_het_=0.14).

**Figure 4:**
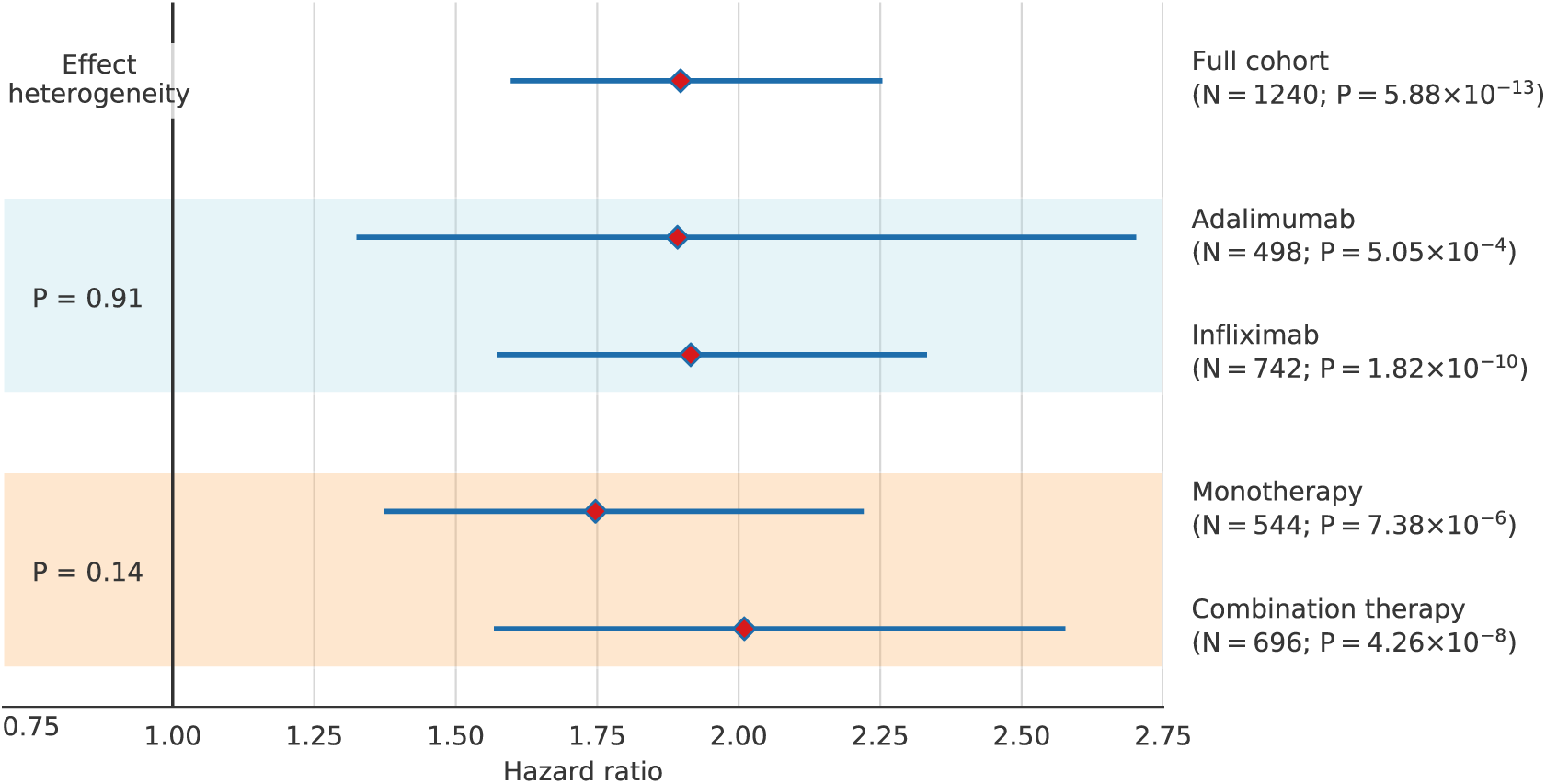
HLA-DQA1*05 has a consistent effect on immunogenicity in different patient subgroups. We repeated the proportional hazard association analysis, separating the full cohort into subgroups by drug and therapy type. Estimated hazard ratios and standard errors between the pairings were compared using a heterogeneity of effects test. None of the pairs had a significant difference in effect (P<0.05), suggesting that the effect of DQA1*05 on immunogenicity is not affected by these clinical covariates.

There was a multiplicative interaction between HLA-DQA1*05 carriage, immunomodulator use and drug type. The highest rates of immunogenicity (92% at 1 year) were observed in patients treated with infliximab monotherapy who carried HLA-DQA1*05 (figure 5).

**Figure 5:**
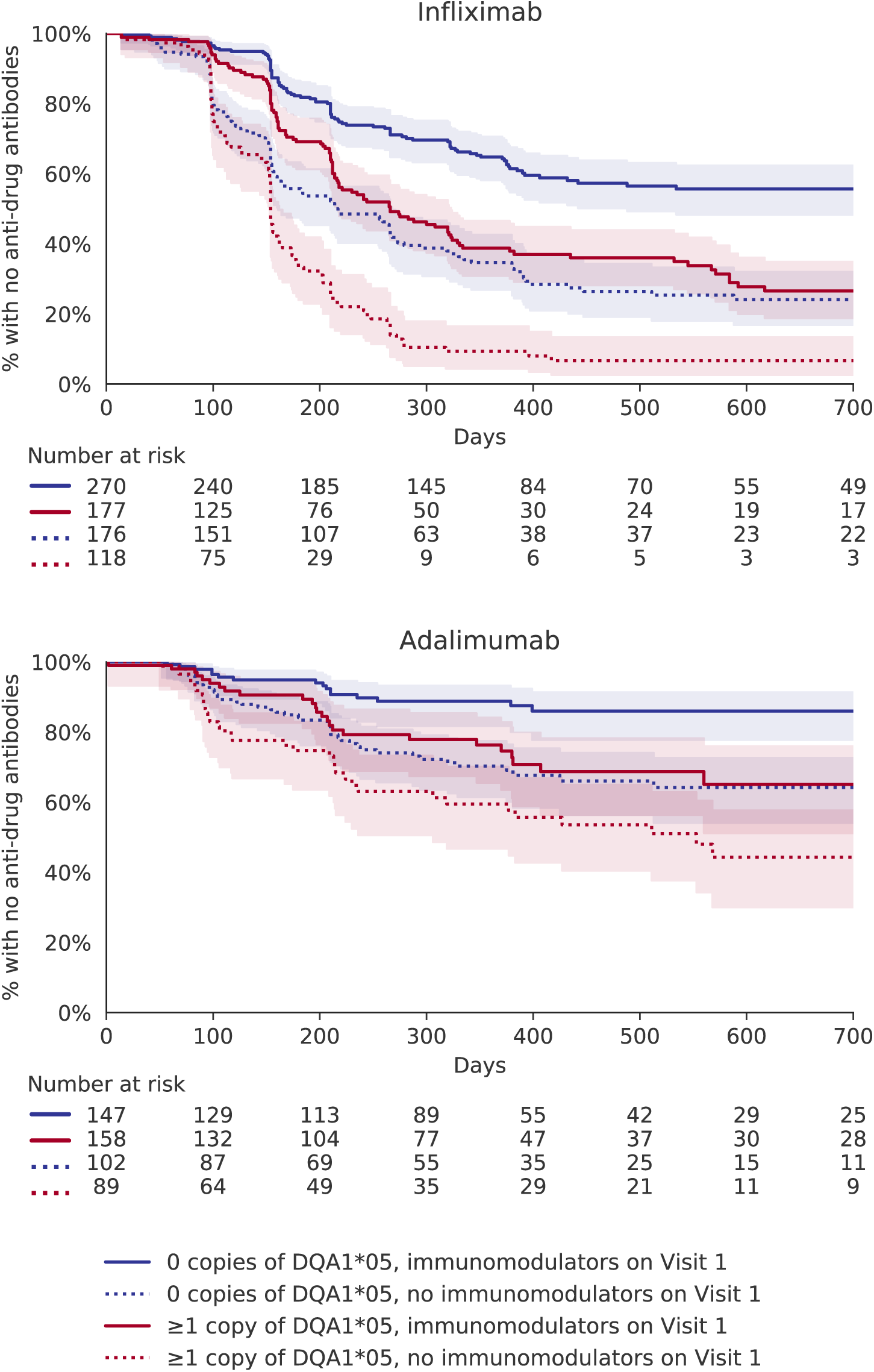
Kaplan–Meier estimator showing the rate of anti-drug antibody development, stratified by the drug, carriage of HLA-DQA1*05 alleles, and treatment regime. Dotted lines indicate patients undergoing anti-TNF monotherapy; solid lines indicate combination therapy with immunomodulators. Red indicates carriers of the HLA-DQA1*05 allele (1 or 2 copies); blue indicates non-carriers. For both drugs and treatment regimes, immunogenicity rates are higher for HLA-DQA1*05 carriers. The X-axis was truncated at 700 days, due to the low number of observations.

## Discussion

We report the first genome-wide significant association with immunogenicity to anti-TNF therapy using the largest prospective cohort study of infliximab and adalimumab in Crohn’s disease. We have demonstrated that carriage of one or more HLA-DQA1*05 alleles confers an almost two-fold risk of immunogenicity to anti-TNF therapy, regardless of concomitant immunomodulator use or drug type (infliximab [Remicade or CT-P13], or adalimumab).

Immunogenicity is more common in patients treated with infliximab than adalimumab. Arguably then, all patients treated with infliximab, as well as HLA-DQA1*05 carriers treated with adalimumab, should also be treated with an immunomodulator. Conversely, patients who do not carry HLA-DQA1*05, in particular those at high risk of opportunistic infections or those with a history of adverse drug reactions to thiopurines and/or methotrexate, could be spared the additional risks of combination therapy and treated with adalimumab monotherapy.

The shared genetic association between HLA-DQA1*05 and immunogenicity to infliximab and adalimumab may explain the widely reported diminishing returns of switching between anti-TNF therapies at loss of response.^29,30^ If the HLA-DQA1*05 immunogenic effect extends to other therapeutic antibodies, then subjects who are HLA-DQA1*05 carriers may be candidates for non-antibody modality therapies such as small molecule drugs.

How variation in the HLA predisposes to immunogenicity is poorly elucidated. The HLA class II gene *HLA-DQA1* is expressed by antigen presenting cells and encodes the alpha chain of the HLA-DQ heterodimer that forms part of the antigen binding site where epitopes are presented to T helper cells. Previous studies have shown that it is possible to map and eliminate potential immunogenic T cell epitopes with the aim of producing a safer, more durable biologic.^31–33^ However, caution needs to be exercised to ensure protein sequence modifications designed to reduce the risk of immunogenicity to patients carrying HLA-DQA1*05 do not put a different group of patients at risk.

We acknowledge several important limitations. Firstly, multiple assays are available to detect anti-drug antibodies and there is no universally accepted, validated threshold to diagnose immunogenicity. Detecting anti-drug antibodies in the presence of drug is difficult and consequently we may have underestimated the proportion of patients with immunogenicity. We deliberately chose a drug tolerant assay in order to mitigate this risk and to minimise the number of false negative patients assigned to the control group, and confirmed that the HLA-DQA1*05 association is robust to the choice of threshold for immunogenicity (appendix figure S5). Secondly, we may have underestimated the contribution of HLA-DQA1*05 to immunogenicity because of the short duration of follow-up in patients who did not continue in the study beyond the first year. Thirdly, because we designed a schedule of visits to minimise patients’ inconvenience, the number of assessments was less for those treated with adalimumab than infliximab. As a result, we might have underestimated rates of immunogenicity amongst adalimumab-treated patients. Finally, unlike the registration trials,^2,34,35^ we did not employ real-time monitoring.

In our final model, which includes HLA-DQA1*05 carriage, drug, and immunomodulator usage (appendix table S2), we have explained 18% of the variance in immunogenicity to anti-TNF in our population. This suggests there are other variables, including other genetic variants and environmental factors, yet to be identified.

Our genome-wide association study was limited to Crohn’s disease patients of European descent. Given that HLA-DQA1*05 is not associated with IBD risk,^36^ the percentage of carriers among our patients (39%) was very similar to that reported in an external British population cohort (38%).^37,38^ As such, we expect HLA-DQA1*05 to have a similar effect on anti-TNF immunogenicity in other patient populations where the frequency of the allele is unchanged. Due to the wide variation in the frequency of HLA-DQA1*05 across ethnic groups^38^, further studies are required to assess the contribution of HLA-DQA1*05 to immunogenicity across populations. Whether HLA-DQA1*05 is also associated with immunogenicity to other biologic drugs also needs to be determined.

In conclusion, immunogenicity is a major concern for patients, regulatory authorities, and the pharmaceutical industry. We have shown that HLA-DQA1*05 identifies patients with IBD at increased risk of immunogenicity regardless of anti-TNF drug. Randomised controlled trials are now needed to determine if pre-treatment genetic testing for HLA-DQA1*05 may help personalise the choice of anti-TNF therapy and allow the targeted use of immunomodulator therapy to minimise risk and maximise response.

## Author contributions

**Concept and design:** T.A., C.A.A., J.C.B., G.H., C.B., T.M., M.P.

**Project manager:** C.B.

**Acquisition, analysis, or interpretation of data:** A.S., D.L.R., L.M., J.C.B., C.A.A.

**Acquisition of replication cohort:** C.L., F.C., M.P., J.M., D.L.R.

**Drafting of the manuscript:** A.S., N.K., L.M., G.H., C.B., D.McG, J.G., C.A.A, T.A.

**Critical revision of the manuscript for important intellectual content:** All authors.

**Statistical analysis**: A.S., L.M., J.C.B., C.A.A.

**Obtained funding:** T.A., C.A.A.

**Administrative, technical, or material support:** All remaining authors contributed by submitting a substantial number of samples in line with ICMJE criteria

### Conflicts of Interest Disclosures

All authors have completed and submitted the ICMJE Form for Disclosure of Potential Conflicts of Interest and declare: G.J.W has consulted for AbbVie and received honoraria from Falk and AbbVie for unrelated topics and a fellowship from NIHR. G.A.H reports non-financial support from AbbVie, outside the submitted work; and that he is now an employee of AbbVie and owns stock in the company. N.A.K has consulted for Falk and received honoraria from Falk, Allergan, Pharmacosmos and Takeda for unrelated topics and is a deputy editor of Alimentary Pharmacology & Therapeutics Journal. J.R.G received honoraria from Falk, Abbvie and Shield therapeutics for unrelated topics. T.A has received unrestricted research grants, advisory board fees, speaker honorariums and support to attend international meetings from AbbVie, Merck, Janssen, Takeda, Ferring, Tillotts, Ferring, Pfizer, NAPP, Celltrion, Hospira for unrelated topics, no financial relationships with any organizations that might have an interest in the submitted work in the previous three years; no other relationships or activities that could appear to have influenced the submitted work.

### Funding/Support

PANTS is an investigator-led study funded by CORE (renamed Guts UK in 2018), the research charity of the British Society of Gastroenterology, and by unrestricted educational grants from Abbvie Inc, USA, Merck Sharp & Dohme Ltd, Ireland, NAPP pharmaceuticals Ltd, UK, Pfizer Ltd, USA, Celltrion Healthcare, South Korea and Cure Crohn’s Colitis (Scottish IBD Charity). The drug and antibody assays were provided at reduced cost by Immundiagnostik AG. Laboratory tests were undertaken by the Exeter Blood Sciences Laboratory at the Royal Devon & Exeter (RD&E) NHS Trust (https://www.exeterlaboratory.com). The sponsor of the PANTS study is the Royal Devon and Exeter NHS Foundation Trust.

AS, LM, DLR, JCB and CAA are funded by the Wellcome Trust, who also funded the genotyping of all samples (206194). TM is funded by an NIHR HEE Senior Lectureship. The views expressed in this publication are those of the author(s) and not necessarily those of the NHS, HEE, NIHR or the Department of Health.

### Role of the Funder/Sponsor

None of the listed funders had a role in the design and conduct of the study; collection, management, analysis, and interpretation of the data; preparation, review, and decision to submit the manuscript for publication.

### Additional Contributions

We would also like to acknowledge the study coordinators of the Exeter IBD Research Group, United Kingdom: Marian Parkinson and Helen Gardner-Thorpe for their ongoing administrative support to the study. We would also like to thank the NIHR UK IBD BioResourse for help establishing the replication cohort.

## Bibliography

1. What’s Trending in Monoclonal Antibodies (Markets by Structure, by Target, and by Indication). Kalorama Information, 2018.

2. Vermeire S, Gils A, Accossato P, Lula S, Marren A. Immunogenicity of biologics in inflammatory bowel disease. Therap Adv Gastroenterol 2018; 11: 1756283X17750355.

3. Mogilevski T, Sparrow MP. Infliximab Versus Adalimumab in Patients with Biologic-Naïve Crohn’s Disease: Is the Difference Real? Dig Dis Sci 2018; 63: 1094–6.

4. Colombel JF, Sandborn WJ, Reinisch W, et al. Infliximab, azathioprine, or combination therapy for Crohn’s disease. N Engl J Med 2010; 362: 1383–95.

5. Panaccione R, Ghosh S, Middleton S, et al. Combination therapy with infliximab and azathioprine is superior to monotherapy with either agent in ulcerative colitis. Gastroenterology 2014; 146: 392–400.e3.

6. D’Haens G, Reinisch W, Panaccione R, et al. Lymphoma Risk and Overall Safety Profile of Adalimumab in Patients With Crohn’s Disease With up to 6 Years of Follow-Up in the Pyramid Registry. Am J Gastroenterol 2018; 113: 872–82.

7. Lichtenstein GR, Feagan BG, Cohen RD, et al. Drug therapies and the risk of malignancy in Crohn’s disease: results from the TREATTM Registry. Am J Gastroenterol 2014; 109: 212–23.

8. Lichtenstein GR, Feagan BG, Cohen RD, et al. Serious infection and mortality in patients with Crohn’s disease: more than 5 years of follow-up in the TREATTM registry. Am J Gastroenterol 2012; 107: 1409–22.

9. Osterman MT, Sandborn WJ, Colombel J-F, et al. Increased risk of malignancy with adalimumab combination therapy, compared with monotherapy, for Crohn’s disease. Gastroenterology 2014; 146: 941–9.

10. Lichtenstein GR, Rutgeerts P, Sandborn WJ, et al. A pooled analysis of infections, malignancy, and mortality in infliximab- and immunomodulator-treated adult patients with inflammatory bowel disease. Am J Gastroenterol 2012; 107: 1051–63.

11. Romero-Cara P, Torres-Moreno D, Pedregosa J, et al. A FCGR3A polymorphism predicts anti-drug antibodies in chronic inflammatory bowel disease patients treated with anti-TNF. Int J Med Sci 2018; 15: 10–5.

12. Billiet T, Vande Casteele N, Van Stappen T, et al. Immunogenicity to infliximab is associated with HLA-DRB1. Gut 2015; 64: 1344–5.

13. Liu M, Degner J, Davis JW, et al. Identification of HLA-DRB1 association to adalimumab immunogenicity. PLoS ONE 2018; 13: e0195325.

14. Perry M, Bewshea C, Brown R, So K, Ahmad T, McDonald T. Infliximab and adalimumab are stable in whole blood clotted samples for seven days at room temperature. Ann Clin Biochem 2015; 52: 672–4.

15. Shah TS, Liu JZ, Floyd JAB, et al. optiCall: a robust genotype-calling algorithm for rare, low-frequency and common variants. Bioinformatics 2012; 28: 1598–603.

16. de Lange KM, Moutsianas L, Lee JC, et al. Genome-wide association study implicates immune activation of multiple integrin genes in inflammatory bowel disease. Nat Genet 2017; 49: 256–61.

17. McCarthy S, Das S, Kretzschmar W, et al. A reference panel of 64,976 haplotypes for genotype imputation. Nat Genet 2016; 48: 1279–83.

18. Zheng X, Shen J, Cox C, et al. HIBAG—HLA genotype imputation with attribute bagging. Pharmacogenomics J 2014; 14: 192–200.

19. Robinson J, Halliwell JA, McWilliam H, Lopez R, Marsh SGE. IPD—the Immuno Polymorphism Database. Nucleic Acids Res 2013; 41: D1234–40.

20. Motyer A, Vukcevic D, Dilthey A, Donnelly P, McVean G, Leslie S. Practical Use of Methods for Imputation of HLA Alleles from SNP Genotype Data. BioRxiv 2016; published online Dec 9. doi:10.1101/091009.

21. Syed H, Jorgensen AL, Morris AP. SurvivalGWAS_SV: software for the analysis of genome-wide association studies of imputed genotypes with “time-to-event” outcomes. BMC Bioinformatics 2017; 18: 265.

22. Davidson-Pilon C, Kalderstam J, Kuhn B, et al. CamDavidsonPilon/lifelines: v0.14.3. 2018; published online May.

23. Therneau TM. A Package for Survival Analysis in S. CRAN, 2015.

24. Viechtbauer W. Conducting meta-analyses in R with the metafor package. Journal of Statistical Software 2010; 36: 1–48.

25. Wellcome Trust Case Control Consortium. Genome-wide association study of 14,000 cases of seven common diseases and 3,000 shared controls. Nature 2007; 447: 661–78.

26. 1000 Genomes Project Consortium, Auton A, Brooks LD, et al. A global reference for human genetic variation. Nature 2015; 526: 68–74.

27. Moutsianas L, Gutierrez-Achury J. Genetic association in the HLA region. Methods Mol Biol 2018; 1793: 111–34.

28. Goncalves J, Santos M, Acurcio R, et al. Antigenic response to CT-P13 and infliximab originator in inflammatory bowel disease patients shows similar epitope recognition. Aliment Pharmacol Ther 2018; published online June 5. doi:10.1111/apt.14808.

29. Panaccione R, Loftus EV, Binion D, et al. Efficacy and safety of adalimumab in Canadian patients with moderate to severe Crohn’s disease: results of the Adalimumab in Canadian SubjeCts with ModErate to Severe Crohn’s DiseaSe (ACCESS) trial. Can J Gastroenterol 2011; 25: 419–25.

30. Sandborn WJ, Rutgeerts P, Enns R, et al. Adalimumab induction therapy for Crohn disease previously treated with infliximab: a randomized trial. Ann Intern Med 2007; 146: 829–38.

31. De Groot AS, Knopp PM, Martin W. De-immunization of therapeutic proteins by T-cell epitope modification. Dev Biol (Basel) 2005; 122: 171–94.

32. De Groot AS, Moise L, McMurry JA, et al. Activation of natural regulatory T cells by IgG Fc-derived peptide “Tregitopes". Blood 2008; 112: 3303–11.

33. Sathish JG, Sethu S, Bielsky M-C, et al. Challenges and approaches for the development of safer immunomodulatory biologics. Nat Rev Drug Discov 2013; 12: 306–24.

34. van der Valk ME, Mangen M-JJ, Severs M, et al. Evolution of Costs of Inflammatory Bowel Disease over Two Years of Follow-Up. PLoS ONE 2016; 11: e0142481.

35. Jørgensen KK, Olsen IC, Goll GL, et al. Switching from originator infliximab to biosimilar CT-P13 compared with maintained treatment with originator infliximab (NOR-SWITCH): a 52-week, randomised, double-blind, non-inferiority trial. Lancet 2017; 389: 2304–16.

36. Goyette P, Boucher G, Mallon D, et al. High-density mapping of the MHC identifies a shared role for HLA-DRB1*01:03 in inflammatory bowel diseases and heterozygous advantage in ulcerative colitis. Nat Genet 2015; 47: 172–9.

37. Thomson W, Barrett JH, Donn R, et al. Juvenile idiopathic arthritis classified by the ILAR criteria: HLA associations in UK patients. Rheumatology (Oxford) 2002; 41: 1183–9.

38. González-Galarza FF, Takeshita LYC, Santos EJM, et al. Allele frequency net 2015 update: new features for HLA epitopes, KIR and disease and HLA adverse drug reaction associations. Nucleic Acids Res 2015; 43: D784–8.

